# Lack of Age-related Respiratory Changes in *Daphnia*

**DOI:** 10.1101/2021.06.05.447222

**Authors:** Cora E. Anderson, Millicent N. Ekwudo, Rachael A Jonas-Closs, Yongmin Cho, Leonid Peshkin, Marc W. Kirschner, Lev Y. Yampolsky

## Abstract

Aging is a multifaceted process of accumulation of damages and waste in cells and tissues; age-related changes in mitochondria and in respiratory metabolism have been in the focus of aging research for decades. Here we investigated age-related changes in respiration rates, lactate/pyruvate ratio, a commonly used proxy for NAD+/NADH balance, and mitochondrial membrane potential in 4 genotypes of an emerging model organism for aging research, a cyclic parthenogen *Daphnia magna*. We show that total body weight-adjusted respiration rate decreases with age, although this decrease is small in magnitude and not observed in anaesthetized animals, thus likely to be accounted for by decrease in locomotion and feeding activity. Lactate/pyruvate ratio and mitochondrial membrane potential (Ψ_mt_) showed no age-related changes, with a possible exception of Ψ_mt_ measured in the optical lobe and in epipodites (excretory organs) in which Ψ_mt_ showed a maximum at middle age. We conclude that actuarial senescence in *Daphnia* is not caused by a decline in respiratory metabolism and discuss possible mechanisms of maintaining mitochondrial healthspan throughout the lifespan.

## Introduction

Ever since - and even somewhat before - the role of mitochondrial membranes in oxidative phosphorylation has been elucidated, researchers asked the question of whether the efficiency of this crucial step of energy metabolism declines with age (Weinbach & Garbus 1956; Gold 1968). Nearly 60 years of research resulted in a substantial body of evidence showing that respiratory metabolism, oxygen consumption and mitochondrial properties are engaged in a complex relationship with age and aging in almost all organisms studied. Membrane phosphorylation is the chief generator of reactive oxygen species, a central factor of damages to membranes, proteins and DNA that accumulate with age. Such damages in turn reduce the intensity of membrane phosphorylation, resulting in reduced oxygen consumption and shifts in the balance of red-ox and phosphorylation reactions, most importantly in NAD+/NADH and ATP/ADP ratios. The decline of NAD+/NADH ratio, in turn, affects NAD+-dependent sirtuin family of protein deacetylases and the PARP family of DNA repair enzymes (Griffiths et al 2020), resulting in imbalance of post-translational regulation of protein activity, decline in autophagy, and impaired DNA repair. In other words, the relationships between aging and respiration are complex and rich in positive and negative feedbacks.

Zooplankton crustacean *Daphnia* is becoming an model organism of choice for aging research due to its ability to reproduce by cyclic parthenogenesis, thus providing an opportunity to test genetically identical individuals in different environments and to generate genetically uniform cohorts (Dudycha JL. 2003; Yampolsky & Galimov 2005; Dudycha & Hassel 2013; Schwartz et al. 2016; Constantinou et al. 2019). Additionally, high permeability of *Daphnia* gut epithelium to a variety of hydrophilic and hydrophobic moieties and high water turnover through the gut results to a relative ease of drug delivery and therefore of pharmacological perturbations in the studies of lifespan and healthspan. However, at the moment, age-related changes in key phenotypes with potential for use as model for biomedical research have not been characterized well (Cho et al., in preparation). Some notable exception are heart rate and locomotion activity shown to markedly decline with age (Constantinou et al. 2019; Cho et al., in preparation), but data on whether these changes are accompanied by decline in respiration rate and redox metabolism are scarce. The fraction of *Daphnia* energy budget allocated to respiration has been shown to remain remarkably stable over the lifespan, with differences observed between genotypes attributable to differences in locomotory activity (Glazier & Calow 1992). Here we report age-related changes (or lack thereof) in active and basal respiration rate, feeding rate, mitochondrial membrane potential and lactate-pyruvate ratio (a proxy for NAD+/NADH ratio) in four clones of *Daphnia magna* in a latitudinal common garden study of individuals of different age from different, uniformly maintained cohorts.

## Materials and Methods

### Daphnia clones, maintenance and cohorts set-up

Four geographically distinct *Daphnia magna* clones previously characterized for life-history and longevity (Coggins et al. 2021; Andershon et al., in preparation) were used in these experiments. Details of geographic origin of these 4 clones and previously measured median lifespans are given in Supplementary Table 1. Stocks and experimental cohorts were maintained at 20 °C in groups of 5 adult females in 100 mL jars with COMBO water (Kilham et al. 1998) under 12:12 photoperiod and fed with *Scenedesmus* culture to the concentration of 100,000 cells/ml daily. Water was replaced and neonates removed every 3 days. This regimen was maintained in the stocks for 2 generations prior to the start of the experiment. Neonates less than 24 h of age were collected from the stock jars and used to establish cohort 1, each next cohort was established from neonates born to the previous cohort 25-35 days apart. Each cohort consisted from 25 individuals per clone, 100 total, maintained in groups of 10-12 in 100 mL of COMBO water until the age of 6 days when they were sexed transferred into new jars in groups of 5 females per 100 mL. Number of surviving individuals was recorded at the time of each water change and individuals from the same clone were combined to maintain the density of 5 individuals per jar. When such pooling was not possible, the volume of water and the amount of food in jars containing less than 5 individuals were adjusted accordingly. Individuals used for experiments were recorded as censored and not returned to the cohorts. Survival data was analyzed using Kaplan-Meier and Proportional Hazards models using JMP Reliability and Survival platform (Ver. 10; SAS Institute 2012). Only the first 3 cohorts were followed to maximal lifespan, so only these cohorts are used in formal survival analysis. Efforts have been made to ensure equal representation of all 4 clones in all ages in all experiments, despite lower portion of cohorts in two out of four clones surviving to the most advanced age; this was not, however, always possible. Specifically, in the rhodamine assays (see below) the two short-lived clones are not represented in the most advanced age classes.

**Table 1.**
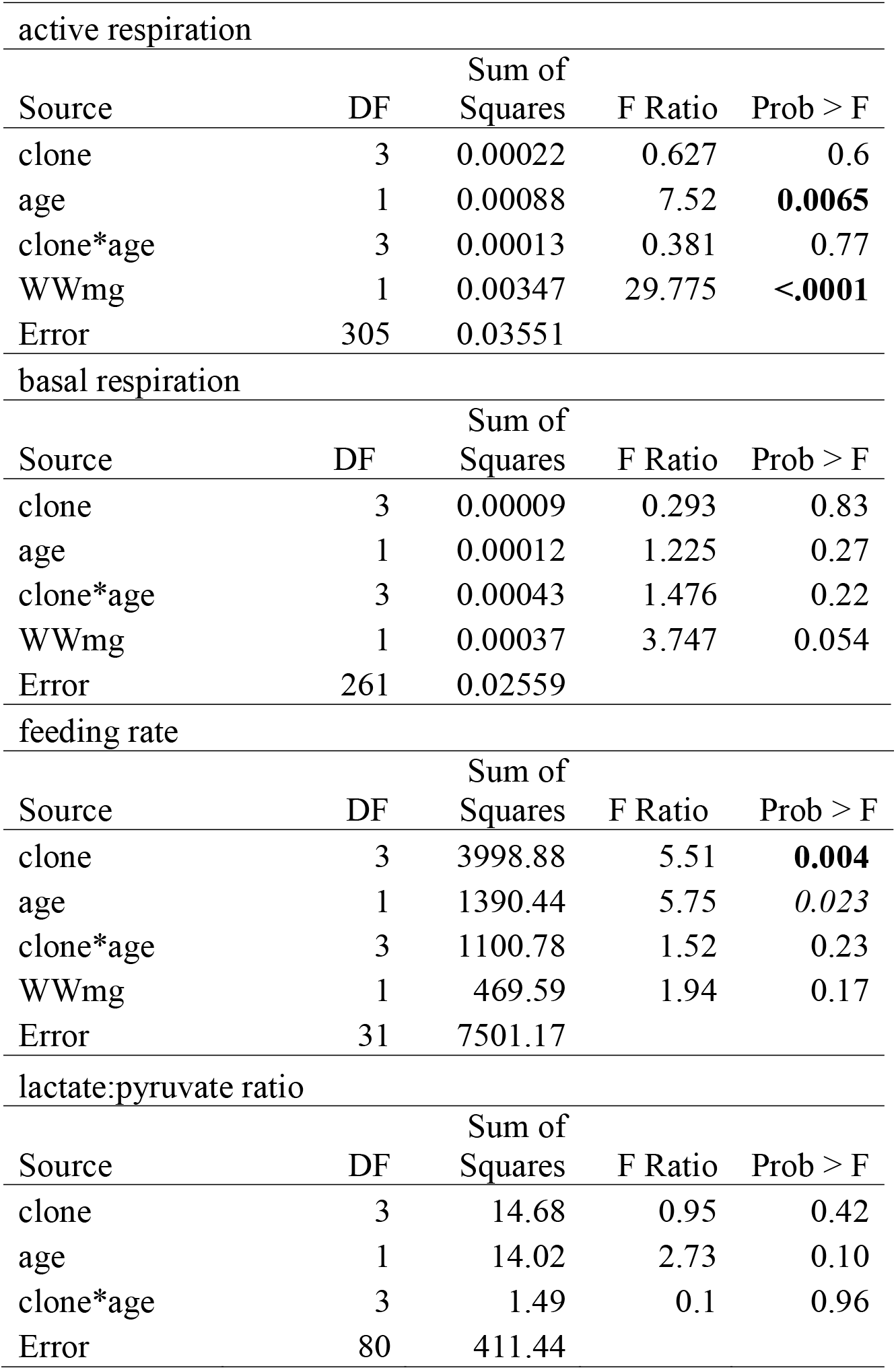
Analysis of variance of the effect of age on active and basal respiration rate (μg O_2_/min/individual) and feeding rate in 4 clones of *D.magna.* Wet weight included in the models as a covariable, except for lactate-pyruvate ratio. P-values < 0.01 shown in **bold**, <0.05 in *italics*.

| active respiration |  |  |  |  |
| --- | --- | --- | --- | --- |
| Source | DF | Sum of Squares | F Ratio | Prob > F |
| clone | 3 | 0.00022 | 0.627 | 0.6 |
| age | 1 | 0.00088 | 7.52 | **0.0065** |
| clone*age | 3 | 0.00013 | 0.381 | 0.77 |
| WWmg | 1 | 0.00347 | 29.775 | **<.0001** |
| Error | 305 | 0.03551 |  |  |
| basal respiration |  |  |  |  |
| Source | DF | Sum of Squares | F Ratio | Prob > F |
| clone | 3 | 0.00009 | 0.293 | 0.83 |
| age | 1 | 0.00012 | 1.225 | 0.27 |
| clone*age | 3 | 0.00043 | 1.476 | 0.22 |
| WWmg | 1 | 0.00037 | 3.747 | 0.054 |
| Error | 261 | 0.02559 |  |  |
| feeding rate |  |  |  |  |
| Source | DF | Sum of Squares | F Ratio | Prob > F |
| clone | 3 | 3998.88 | 5.51 | **0.004** |
| age | 1 | 1390.44 | 5.75 | *0.023* |
| clone*age | 3 | 1100.78 | 1.52 | 0.23 |
| WWmg | 1 | 469.59 | 1.94 | 0.17 |
| Error | 31 | 7501.17 |  |  |
| lactate:pyruvate ratio |  |  |  |  |
| Source | DF | Sum of Squares | F Ratio | Prob > F |
| clone | 3 | 14.68 | 0.95 | 0.42 |
| age | 1 | 14.02 | 2.73 | 0.10 |
| clone*age | 3 | 1.49 | 0.1 | 0.96 |
| Error | 80 | 411.44 |  |  |

### Respiration rates

Respiration rates were measured using Loligo respirometer (Loligo® Systems, Viborg, Denmark) with either 1700 μL well (for active metabolism) or 200 μL well plates. Each plate included 3 blanks. Single *Daphnia* were placed individually into the wells containing either 1700 μL of COMBO water or 200 μL 1% urethane solution in COMBO water (Philippova & Postnov 1988) for active and basal metabolism, respectively, with water, plates, and sensors equilibrated to 20 °C. Oxygen concentrations were measured every 15 seconds after initial 15 minute break-in period for 45 minutes or until the concentration decreased by 2 mg/L from the blank readings. Oxygen consumption per minute was calculated from a linear regression of oxygen concentration over time; mean blank estimates subtracted. After measurements *Daphnia* were removed from Loligo wells, their body length and wet weight were measured, and individuals were frozen for lactate and puruvate determination. Respiration rates were normalized by individuals’ wet weight only for presentation purposes. To detect a significant change across ages, however, respiration rates were calculated per individual and wet weight was used as a co-variable. Same approach was used for the analysis of feeding rate.

### Feeding rate

Feeding (filtering) rate was determined by the decrease of chlorophyll fluorescence using Loligo respirometers equipped with regular plastic 24-well plates, each well containing a single *Daphnia* and COMBO water initially containing *Scenedesmus* algae with the concentration of 200,000 cells/mL. Measurements were done in the initial algae suspension and after 18 hours of feeding in a dark incubator at 20 °C and the amount consumed was determined from the difference in fluorescence using a calibration with a linear range between 25,000 and 400,000 cells/mL. After measurements *Daphnia* wet weight was determined to be used for normalization purposes and individuals were placed back into their corresponding cohort jars.

### Lactate/pyruvate ratio

To quantify red-ox equilibrium whole body extract lactate to pyruvate ratio was measured in *Daphnia* of different age. Lactate to pyruvate ratio is commonly used as a proxy to cytosolic NAD+ to NADH ratio, as unambiguous determination of NAD+ and NADH is difficult due to larger portion of these coenzymes being bound to proteins and both substrates of lactate dehydrogenase are at an equilibrium with the reduced and oxidized forms of the coenzyme (Williamson et al. 1967; Mintun et al 2004). The estimated NAD+/NADH ratio is inversely proportional to the lactate/pyruvate ratio, with the coefficient of proportionality being the inverse of the equilibrium constant of the lactate dehydrogenase reaction. This approach is not problem-free and deviations from the equilibrium may result in biased estimates of NAD+/NADH balance (Sun et al. 2012), but it is conventional as a measure of red-ox balance changes over age or in diagnostics of various clinical conditions related to mitochondrial health and hypoxia (Debray et al 2007; Rimachi et al. 2012).

Lactate and pyruvate concentrations in whole body were measured using CellBiolab lactate (MET-5013) and pyruvate (MET-5029) fluorometric assay kits. The same individuals used in respiration measurements were weighed to determine wet weight and frozen at −80°C. Each Daphnia was homogenized in 100 μL ice-cold PBS with a pestle and the homogenates were centrifuged at 4°C for 4 min. 25 μL of supernatant were pipetted into each of the lactate and pyruvate assay plates with well contents according to manufacturer’s protocols. Additionally, 25 μL of the supernatant was used to quantify soluble proteins by Bradford assay.

### Mitochondrial membrane potential

Individual *Daphnia* were placed in 0.5 mL of COMBO water containing 4 μM solution of rhodamine123 (made from 1000X stock in DMSO) in the dark for 24 hours. After 3-fold rinsing with COMBO water individuals were photographed using a Leica DM3000 fluorescent microscope with a 10x objective (0.22 aperture) equipped with Leica DFc450C camera using the 488 nm excitation / broadband (>515 nm) emission filter. The following ROI were selected peduncle of antenna-2 (striated muscle), heart, brain, optical lobe, 2nd epipodite or “gills” (thoracic appendage branch with osmoregulatory functions), and non-neural (parenhymal) head tissue. Median fluorescence (background subtracted) was recorded with exposure of 100 ms with gain 1. The concentration of 4 μM rhodamine123 was chosen as being close to Michaelis-Menten constant of the process of accumulation of the dye in mitochondria (Hasan, 2018; Anderson et al. in preparation). Average values of 2-3 replicate individuals from the same clone and the same age measured on the same date are reported.

## Results

### Cohorts

As expected, the two short-lived and the two long-lived clones differed from each other in lifespan, although the very old individuals from the short-lived FI tended to survive the longest (Supplementary Figure S1). The difference among clones, however, was not consistent across cohorts, as the clone effect disappears and a weak cloneXcohort interaction is detected (Supplementary Table 2). By the highest age studied *Daphnia* in all clones experience actuarial aging with mortality increasing approximately 7-fold relative to initial mortality (Supplementary Figure S1B).

### Respiration rates

Wet weight-normalized respiration rate decreased linearly with age, both in active and in anesthetised *Daphnia* (Fig.1). However, the decline of respiration per individual was statistically significant, with wet weight as a co-variable, only for active metabolism measurements (Table 1). There was no evidence of clone-by-age interaction for either normalization. There were no correlation between respiration rate and either the number of eggs in the clutch carried at the time of measurement or with the development stage of the clutch. There was, however, a slight difference active respiration rate between individuals carrying a clutch as compared to those carrying no clutch and without visible ovaries or those carrying no clutch but with visible ovaries (i.e. within 24 hours of producing a new clutch). This difference remained significant when both wet weight and age were accounted for by including these terms as co-variables. It is unlikely that this difference indicates disproportionally higher respiration rate of embryos, as it was only observed in actively moving individuals. Rather, it may indicate higher expenditures on active swimming while carrying a clutch.

**Fig. 1.**
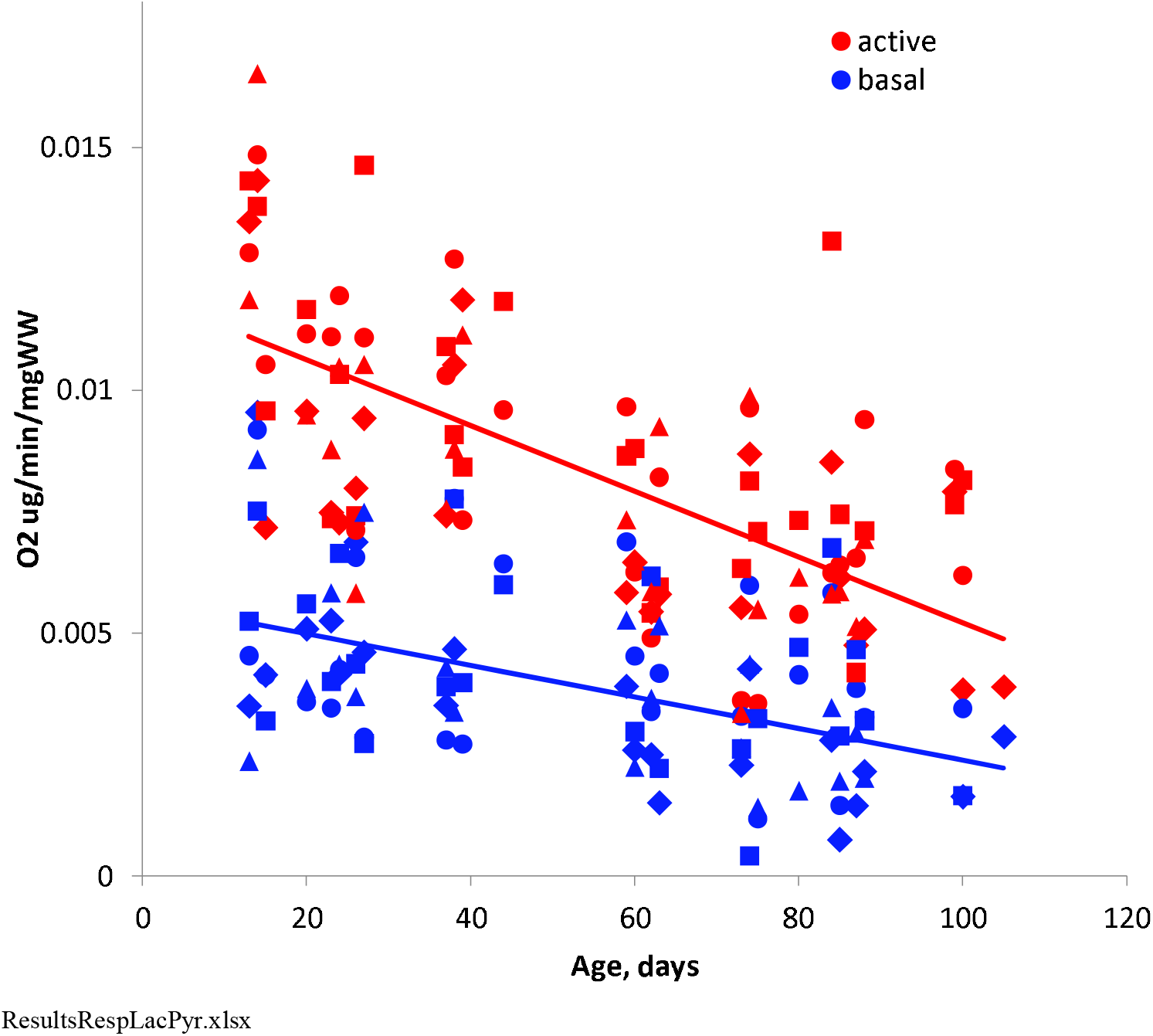
Active (red) and basal (blue) respiration rate in four *Daphnia* clones as a function of age. Each dot represents average of 2-4 replicate individuals measured at a given day. Regression lines drawn through all replicates. See Table 1 for statistical tests. Clones 2-letter codes: FI - diamonds, GB - circles, HU - squares, IL - triangles.

### Feeding rate

There was a consistent decline in feeding rate normalized by wet weight with individual’s age (Fig. 2). This decline is not a normalization artifact, as the age term remains significant with wet weight used as a covariable (Table 1). There was also a significant clone effect with different clones showing different feeding rate with body size accounted for.

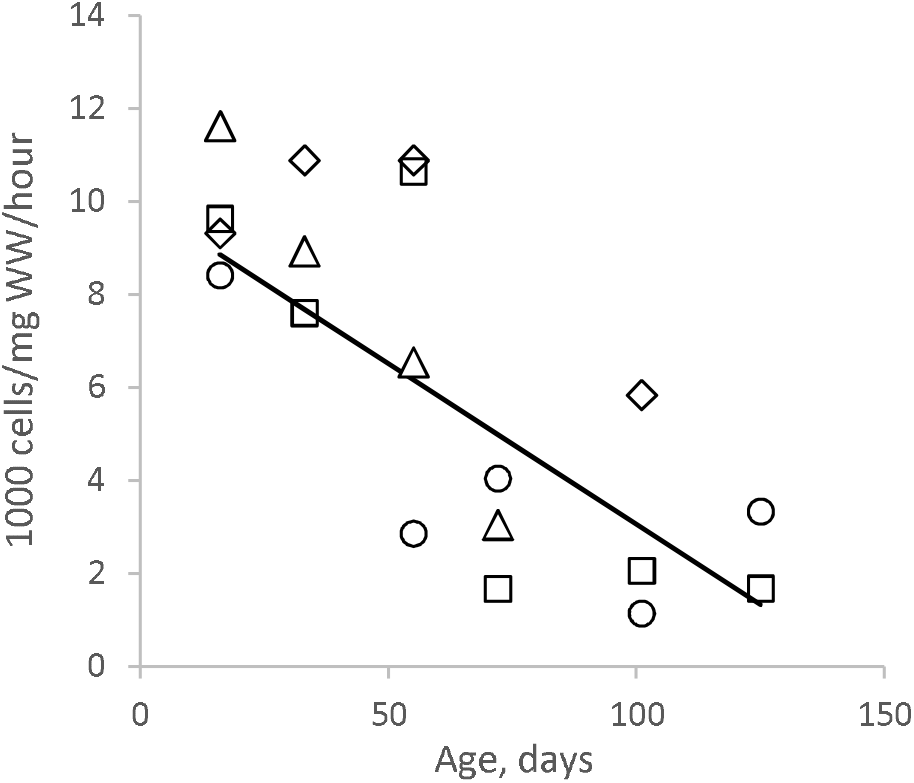

### Lactate/pyruvate ratio

Overall lactate/pyruvate ratio was found to be low: 3.70 ±(SE) 0.23. Contrary to the predictions, lactate/pyruvate ratio did not increase with age. If anything, there was a slight decrease, not reaching a statistical significance and less than 25% relative change over the entire lifespan in magnitude (Fig.3, Table 1). The slight decrease was caused by equally slight increase in pyruvate concentration with completely constant lactate concentration (both normalized by protein content) over age (data not shown).

**Fig. 3.**
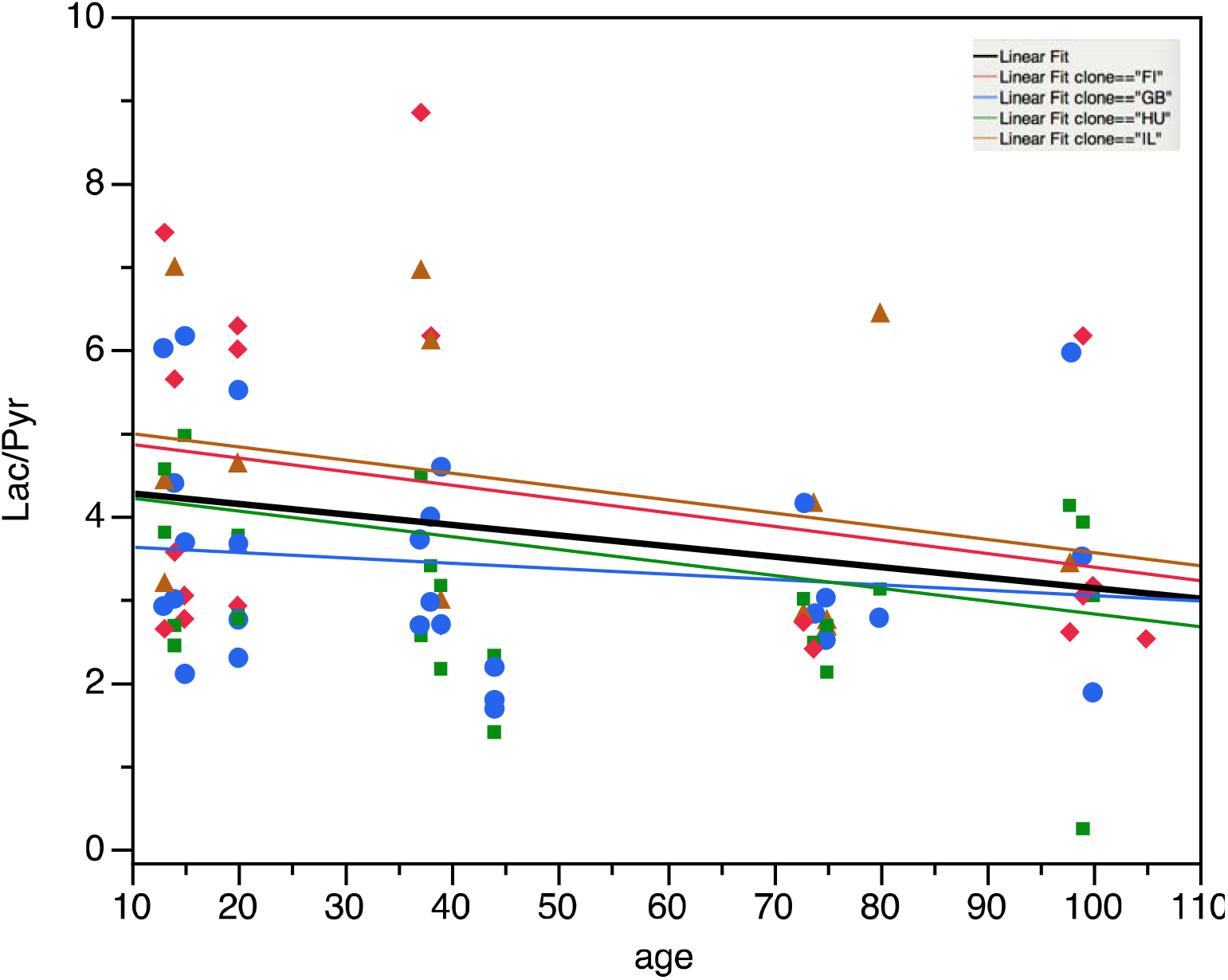
No significant changes in lactate/pyruvate ratio with age (days) in 4 *Daphnia* clones. Overall regression coefficient R = −0.0126 ± 0.0077. Symbols as on Fig.1.

### Mitochondrial membrane potential

Rhodamin-123 assay of mitochondrial membrane potential (Ψ_mt_) showed little change with individuals’ age in most organs and tissues studied (Fig. 4, Table 2, Supplementary Table S3). The only exceptions were the optical lobe (Fig. 4 D), in which there seemed to be a maximum of membrane potential around the age of 75-80 days with a slight decline afterwards (the quadratic term of a second degree polynomial regression R=−0.00223±0.00095; P<0.0254) and the epipodite (gill) tissue in which Ψ_mt_ showed a consistent decline with age (Fig. 4E). Such differential effect of age in different tissues (ROIs) resulted in a significant age*ROI interaction term (Table 2).

**Fig. 4.**
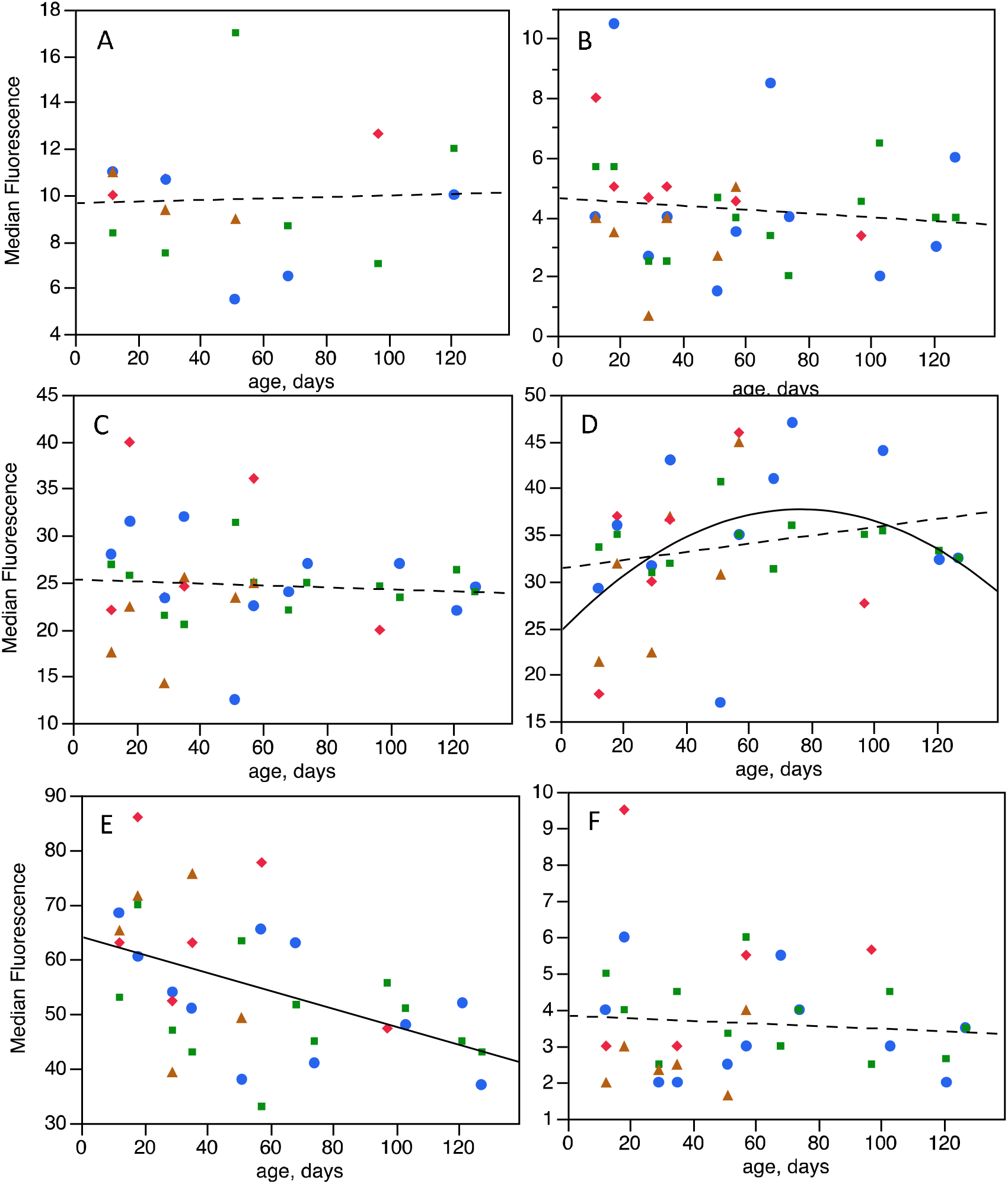
Rhodamine-123 fluorescence (median, background-subtracted) in various tissue of *Daphnia* as a function of age. High values are indicative of higher mitochondrial membrane potential. Averages of 2-4 replicates per clone per age measured on the same data are shown. A: antenna-2 (striated muscle); B: heart; C: brain; D: optical lobe; E: epipodite; F: non-neural head tissue. Regressions shown: dashed line where not significant, solid lines where significant (P< 0.005), quadratic regression shown where the 2nd degree term significant (P<0.025). Symbols as on previous figures.

**Table 2.** Analysis of variance of the effects of age, clone, and tissue (region of interest, ROI) on rhodamine-123 fluorescence. ROI’s studied include: muscle tissues (antenna-2, heart), neural tissue (brain, optical lobe), non-neural head tissue and epipodite (gill). See Fig. 4 and Supplementary table S3 for the analysis for each ROI separately.

| Source | DF | Sum of Squares | F Ratio | Prob > F |
| --- | --- | --- | --- | --- |
| Clone | 3 | 206.09 | 0.94 | 0.42 |
| age | 1 | 115.47 | 1.59 | 0.21 |
| Clone*age | 3 | 51.31 | 0.23 | 0.87 |
| ROI | 5 | 77296.49 | 212.26 | <.0001 |
| Clone*ROI | 15 | 615.98 | 0.56 | 0.9 |
| age*ROI | 5 | 1694.01 | 4.65 | 0.0004 |
| Clone*age*ROI | 15 | 706.37 | 0.65 | 0.84 |
| Error | 387 | 28185.84 |  |  |

## Discussion

We measured several parameters of *Daphnia* respiratory red-ox metabolism across a wide range of ages, including highly advanced age when actuarial senescence is very apparent and to which only ~10% of the initial cohort survives. We did not observe any significant changes in respiration rate, lactate and pyruvate concentrations and their ratio, and, with one exception, Ψ_mt_, that cannot be accounted for by age-related decrease of locomotory activity (well documented previously) and/or by the decrease of filtering activity as described here. Thus, *Daphnia* is capable to maintain active aerobic respiration even in the part of the lifespan when many other organs fail and the age is approaching maximal lifespan.

We observed that while the total respiration rate declined with age, the basal rate measured in urethane-anesthetized animals did not. Urethane reversibly paralyzes striated muscles, in *Daphnia* resulting in ceased antennal beat and thoracic appendages movement. Both result in decreased oxygen demand, but the immobilization of thoracic appendages also effects the supply of oxygen. Although epipodites are not an exclusive site of dissolved gases exchange with the environment (Pirow et al. 1999; Smirnov 2017), the circulation of water created by the movement of thoracic limbs is critical for oxygen supply. In immobilized animals oxygen enters tissues only through diffusion. We believe that it is primarily the immobilization of these appendages that created a lower, and age-independent level of oxygen consumption in urethane-treated animals. It is unlikely that immobilizing antennae could have had a profound effect on respiratory demand in our experimental set-up, because not much swimming occurs in the 1.7 mL, 12 mm diameter Loligo wells even in untreated individuals, where their locomotion is likely to be suppressed by frequent encounters with the well walls. Post-measurement observations showed that these *Daphnia* were invariably found at the top of the well, possibly avoiding the excitation light coming from the bottom of the well, showing no swimming activity. However, it is impossible, in this experiment, to distinguish between reduced supply and reduced demand effects of immobilization. Because the movement of thoracic appendages is also required for filter-feeding, the observed decline in feeding rate with age should also result in both reduced demand and reduced supply of oxygen. It is probably not by chance that the only tissue in which we observed a decline of Ψ_mt_ with age was the epipodites, or “gills” - leaf-shaped leaf-shaped branches of thoracic limbs in cladocerans. These mitochondria-rich organs (Kikuchi 1983) have the function of dissolved gasses and ions exchange with the environment and osmoregulation in the adult *Daphnia* (Pirow et al. 1999; Smirnov 2017).

Even keeping in mind that members of all clonal cohorts are genetic copies of each other (safe for de novo mutations) and shared current, maternal and grandmaternal environments, thus making them likely to be epigenetically uniform, it is impossible to fully exclude cohort heterogeneity. It is entirely possible that even initially identical individuals may diverge in their pathways of accumulating damages throughout the lifespan. It would be necessary to conduct a longitudinal experiment with individually maintained *Daphnia,* tracking individual changes in respiration rate and correlating them with lifespan. Such a study will have a number of limitations: lactate/pyruvate ratio could only be measured in embryos or neonates rather than maternal tissue, while the rhodamine assays may have an effect on survival and/or further assays in the same individual (it does not cause immediate mortality if the specimen is exposed to the excitation light for less than a minute; L.Yampolsky, unpublished). Yet, such an experiment might reveal a decrease in respiration immediately prior to death. If such decreases become more apparent with age, this finding might refute the conclusion of the present study. It would then interesting to investigate what genetic or epigenetic changes or damages cause such divergence of respiratory health in identical individuals.

However, until such a longitudinal study is conducted, our conclusion stands and it raises the question about its generality. Is maintenance of respiratory metabolism and mitochondrial membrane potential throughout the lifespan a *Daphnia-*specific phenomenon? Can it be used to pinpoint potential maintenance investments and/or rejuvenation mechanism that *Daphnia* employs to preserve mitochondrial function and red-ox balance all the way into very advanced age? Is the observed decrease in total oxygen consumption with age fully explainable by the reduced decline in feeding rate and associated decline in thoracic limbs movement and is such decline operating through decreased demand or decreased supply of oxygen? More generally, it is not clear whether the observed decrease in total respiration rate is a result of age-related damages (i.e., reduction of mitochondrial efficiency) or of age-related decrease in demand. The observed decrease in Ψ_mt_ in the very organ responsible for oxygen delivery to tissues is consistent with both pathways.

## Conclusions

Contrary to expectations we did not detect any internal, motility-independent age-related changes in respiration rate, lactate/pyruvate ratio and mitochondrial membrane potential (Ψ_mt_) in *Daphnia.* The only possible exception is the decrease of Ψ_mt_ in epipodites (gills) - thoracic limbs branch responsible for gas and ion exchange with the environment, indicative of either reduced demand or reduced supply of oxygen in older *Daphnia.*

## Acknowledgements

We are grateful to Loligo Systerms personnel for advise on setup of Loligo respirometry, to Morad Malek for laboratory assistance and to D.Kumar and G. Arceo-Gomez for a access to laboratory equipment.

