## Supplementary Tables and Figures for "Lack of Age-related Respiratory Changes in *Daphnia*"

A

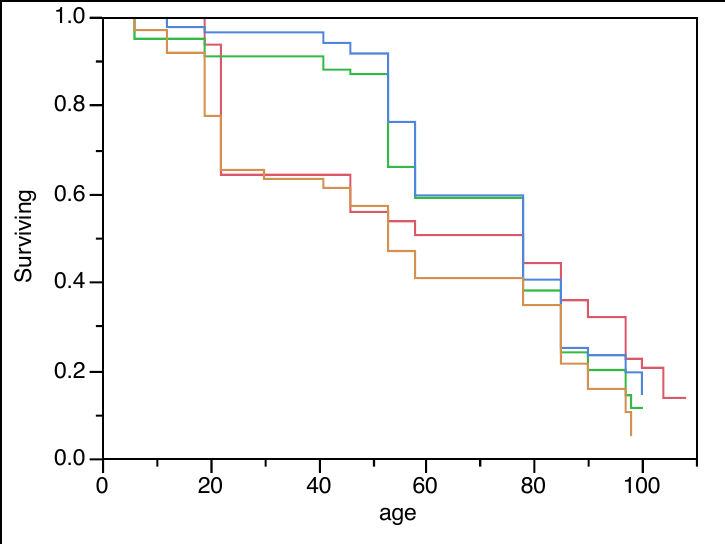

IL	FI

HU	GB

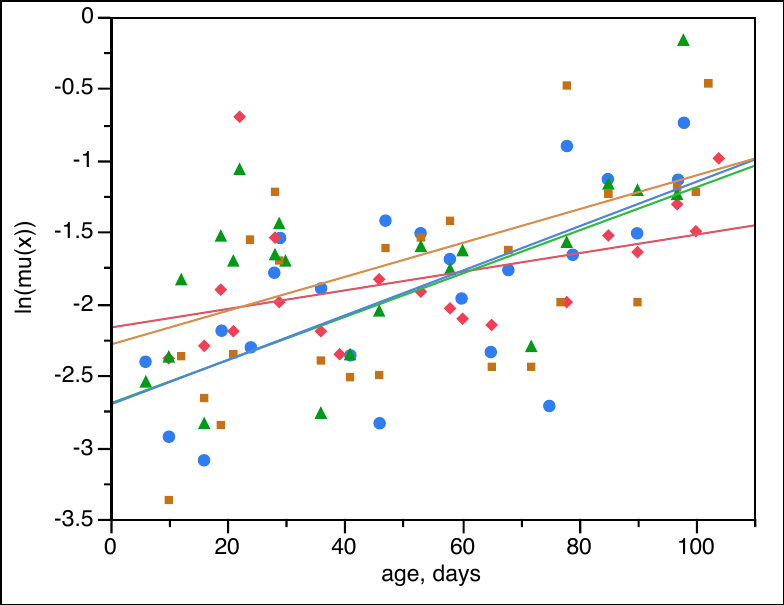

B

Fig. S1. A. Survival of 4 clones in cohort 1. Log-rank test for differences in lifespan among clones: X^2^=9.34, d.f.=3, P<0.026; Wilcoxon test X^2^=14.1, d.f.=3, P<0.003. Clones' 2-letter IDs shown. B. Gompertz model (natural logarithm of mortality rate as a linear function of age) fitted to each clone separately.

O2 ug/min/mgWW

|  | active | basal |
| --- | --- | --- |
| no clutch, no ovaries | 0.0228082 | 0.0104265 |
| no clutch, ovaries | 0.0256141 | 0.0113262 |
| clutch | 0.0277497 | 0.012833 |

Fig. S2. Active and basal respiration rate in individuals carrying no clutch, carrying no clutch, but about to produce one (visible ovaries) and carrying a clutch of eggs (least mean square means with wet weight and age accounted for). Active metabolism: F=4.192, P<0.016, d.f.'s 2, 300. Basal metabolism: F=1.041, P>0.35; d.f.'s 2, 254. Tukey test (P=0.05) pairwise comparison shown (separately for active and basal metabolism data).

Table S1 Location and habitat type of origin of Daphnia genotypes used. First two letters of Clone ID are used to identify clones throughout the paper.

*Anderson et al., in preparation (Experiment 1).

| **CloneID** | **Type of Habitat** | **Latitude** | **Longitude** | **Median lifespan, days ***  **(95% CI)** |
| --- | --- | --- | --- | --- |
| FI-FSP1-16-2 | summer rock pool | 60° 10.062" | 25° 47.677" | 36 (22-42) |
| GB-EL75-69 | year-round pond | 51°30′26″ | -0°7′39″ | 64 (62-68) |
| HU-K-6 | lake | 46° 47' 33.3" | 19° 10' 53.84" | 72 (67-77) |
| IL-MI-8 | Mediterranean pond | 31° 42' 52.42" | 35° 3' 3.38" | 54 (46-57) |

Table S2. Proportional hazards test of the effects of clones and cohorts (cohorts 1-3) on age-specific survival in 4 *Daphnia* clones.

| Source | DF | L-R ChiSquare | Prob>ChiSq |
| --- | --- | --- | --- |
| clone | 3 | 1.772 | 0.62 |
| cohort | 2 | 6.153 | 0.05 |
| clone*cohort | 6 | 14.317 | 0.03 |

Table S3. Analysis of variance of the effects of clones and age on rhodamine-123 fluorescence by ROI (tissue)

| ROI: antenna-2 | |  |  |  |
| --- | --- | --- | --- | --- |
| Source | DF | SS | F | P |
| Clone | 3 | 26.61 | 0.48 | 0.70 |
| age | 1 | 0.61 | 0.03 | 0.86 |
| Clone*age | 3 | 22.88 | 0.42 | 0.74 |
| Error | 39 | 714.21 |  |  |
| ROI: brain |  |  |  |  |
| Clone | 3 | 27.07 | 0.18 | 0.91 |
| age | 1 | 1 | 0.02 | 0.89 |
| Clone*age | 3 | 151.74 | 1 | 0.40 |
| Error | 70 | 3539.52 |  |  |
| ROI: epipodite | |  |  |  |
| Clone | 3 | 658.16 | 0.96 | 0.42 |
| age | 1 | 1483.51 | 6.5 | 0.013 |
| Clone*age | 3 | 308.9 | 0.45 | 0.72 |
| Error | 69 | 15741.02 |  |  |
| ROI: heart |  |  |  |  |
| Clone | 3 | 12.73 | 0.52 | 0.67 |
| age | 1 | 5.43 | 0.67 | 0.42 |
| Clone*age | 3 | 10.77 | 0.44 | 0.72 |
| Error | 69 | 560.61 |  |  |
| ROI: non-neural head tissue | | |  |  |
| Clone | 3 | 24.11 | 1.75 | 0.16 |
| age | 1 | 0.03 | 0.01 | 0.94 |
| Clone*age | 3 | 6.81 | 0.49 | 0.69 |
| Error | 70 | 321.29 |  |  |
| ROI: optical lobe | |  |  |  |
| Clone | 3 | 56.83 | 0.18 | 0.91 |
| age | 1 | 280.99 | 2.69 | 0.11 |
| Clone*age | 3 | 276.82 | 0.88 | 0.45 |
| Error | 70 | 7309.19 |  |  |
